# Structural basis for transcription activation by Crl through tethering of σ^S^ and RNA polymerase

**DOI:** 10.1101/726851

**Authors:** Alexis Jaramillo Cartagena, Amy B. Banta, Nikhil Sathyan, Wilma Ross, Richard L. Gourse, Elizabeth A. Campbell, Seth A. Darst

## Abstract

In bacteria, a primary σ factor associates with the core RNA polymerase (RNAP) to control most transcription initiation, while alternative σ factors are used to coordinate expression of additional regulons in response to environmental conditions. Many alternative σ factors are negatively regulated by anti-σ factors. In *Escherichia coli, Salmonella enterica*, and many other *γ*-proteobacteria, the transcription factor Crl positively regulates the alternative σ^S^ regulon by promoting the association of σ^S^ with RNAP without interacting with promoter DNA. The molecular mechanism for Crl activity is unknown. Here, we determined a single-particle cryo-electron microscopy structure of Crl-σ^S^-RNAP in an open promoter complex with a σ^S^ regulon promoter. In addition to previously predicted interactions between Crl and domain 2 of σ^S^ (σ^S^), the structure, along with p-benzoylphenylalanine crosslinking, reveals that Crl interacts with a structural element of the RNAP β’ subunit we call the β’-clamp-toe (β’CT). Deletion of the β’CT decreases activation by Crl without affecting basal transcription, highlighting the functional importance of the Crl-β’CT interaction. We conclude that Crl activates σ^S^-dependent transcription in part through stabilizing σ^S^-RNAP by tethering σ^S^ and the β’CT. We propose that Crl, and other transcription activators that may use similar mechanisms, be designated σ-activators.

**Significance Statement:** In bacteria, multiple σ factors can bind to a common core RNA polymerase (RNAP) to alter global transcriptional programs in response to environmental stresses. Many γ-proteobacteria, including the pathogens *Yersinia pestis, Vibrio cholera, Escherichia coli*, and *Salmonella typhimurium*, encode Crl, a transcription factor that activates σ^S^-dependent genes. Many of these genes are involved in processes important for infection, such as biofilm formation. We determined a high-resolution cryo-electron microscopy structure of a Crl-σ^S^-RNAP transcription initiation complex. The structure, combined with biochemical experiments, shows that Crl stabilizes σ^S^-RNAP by tethering σ^S^ directly to the RNAP.

---

Bacterial transcription initiation requires the assembly of a promoter-specificity sigma (σ) factor with the RNA polymerase (RNAP) catalytic core (E, subunit composition α_2_ββ’ω), forming the RNAP holoenzyme (Eσ) (1). Multiple σ factors compete for binding to core RNAP, with each σ factor directing transcription of a specific set of promoters, or regulon (2). An essential primary σ directs most transcription during normal growth conditions, while alternative σ’s direct transcription of regulons in response to metabolic, developmental, and environmental signals (2).

The vast majority of σ factors belong to the σ^70^-family (3), which minimally contain two flexibly linked, conserved structural domains, σ_2_ and σ_4_. In the absence of core RNAP, many σ^70^-family factors have been proposed to adopt a compact conformation where the promoter DNA-binding determinants in σ_2_ (recognizing the promoter −10 element) and σ_4_ (recognizing the promoter −35 element) are inaccessible, explaining why σ^70^-family members bind their cognate promoter sequences very poorly or not at all without the core RNAP. In the holoenzyme, σ factors adopt an open conformation where the σ_2_ and σ_4_ domains are displayed on the RNAP surface with the proper spacing to recognize the −10 and −35 promoter elements, centered 75 to 80 Å apart (4-11).

A key mechanism to control transcription initiation in bacteria is to regulate access of σ factors to the core RNAP with anti-σ factors (12). Anti-σ factors stabilize occlusive inter-domain interactions within the σ factor and/or physically occlude the RNAP interacting surface (13-19). Upon relief of inhibition, the RNAP binding surfaces of the σ factor are exposed, allowing interactions with RNAP.

*Escherichia coli* (*Eco*) has seven σ factors; σ^70^ is the primary (housekeeping) σ, while σ^S^ (encoded by *rpoS*) is the master regulator of transcription programs in the stationary phase of growth as well as in response to various stresses including antibiotics, UV light, low temperature, osmolarity changes, acidity changes, and nutrient depletion (20). In certain conditions, the rapid and efficient expression of genes under σ^S^ control is critical for the survival of bacteria. However, once conditions become favorable for growth the σ^S^ transcription program must be shut down for optimal fitness. For these reasons, the expression of σ^S^ is highly regulated at transcriptional, translational, and post-translational levels (21).

Transcription from σ^S^-dependent promoters can be limited by the Eσ^S^ concentration. To form Eσ^S^, σ^S^ must compete against other σ factors to assemble with free core RNAP, for which σ^S^ has the lowest binding affinity (22). Crl is an ∼16 kDa protein, widely distributed in *γ*-proteobacteria, that specifically activates Eσ^S^ transcription (23, 24). Crl does not bind DNA like most transcription factors (25) but rather acts by directly binding domain 2 of σ^S^ (σ^S^_2_) (26) and stimulating expression of stress response genes, genes required for formation of amyloid curli fibers involved in adhesion and biofilm formation (23), and many other genes in the σ^S^ regulon (24).

Crl accumulates during bacterial exponential growth and reaches peak levels as bacteria enter stationary phase, with levels dropping as cells progress into late stationary phase (27, 28). By contrast, σ^S^ is not detectable until bacteria begin to enter stationary phase, and the level of σ^S^ continues to increase until late stationary phase (29). This interplay in the levels of Crl and σ^S^ suggests a critical role for Crl when the levels of σ^S^ are very low. This is consistent with *in vitro* experiments demonstrating that transcription activation by Crl is most pronounced when σ^S^ concentrations are lowest (30-32). These findings have led to proposals that Crl functions by facilitating the assembly of core RNAP and σ^S^ into Eσ^S^ (26, 30).

Previous studies have determined structures of Eσ^S^ (33) and Crl (34, 35) in isolation. However, understanding the molecular mechanism of Crl has been hindered by the lack of structures of complexes of Crl with σ^S^ or with Eσ^S^. Here, we employed single-particle cryo-electron microscopy (cryo-EM) to determine the structure of Crl bound to an Eσ^S^ open promoter complex (RPo) containing a σ^S^-regulon promoter. Our analysis of the structure, combined with biochemical assays, shows that Crl simultaneously interacts with σ^S^ and core RNAP in the complex, stabilizing Eσ^S^ by tethering σ^S^ with RNAP. We propose that Crl, and other unconventional transcription activators that use a similar mechanism, be designated as σ-activators.

## Results

### *Salmonella* Crl-σ^S^ activates *Eco* core RNAP

For our structural and functional analyses, we studied a complex between *Salmonella enterica* serovar Typhimurium (*Sty*) Crl, *Sty* σ^S^, and *Eco* core RNAP lacking the α C-terminal domains (hereafter designated Crl-Eσ^S^) rather than from the same bacterium because overexpressed *Eco* Crl and *Eco* σ^S^ tended to form insoluble aggregates. *Sty* and *Eco* Crl-σ^S^ have 95% sequence identity over 463 residues. The entire 443 kDa Crl-Eσ^S^ complex has 98.3% sequence identity over 4,576 residues (*SI Appendix*, Table S1). Purified *Sty* Crl activated Eσ^S^ transcription ∼5-fold (compared to no Crl) in an *in vitro* abortive initiation assay on a linear fragment of a σ^S^-regulon promoter, *dps* (36, 37)(Fig. 1A), indicating that our Crl-Eσ^S^ complex is structurally and functionally relevant.

**Fig. 1.**
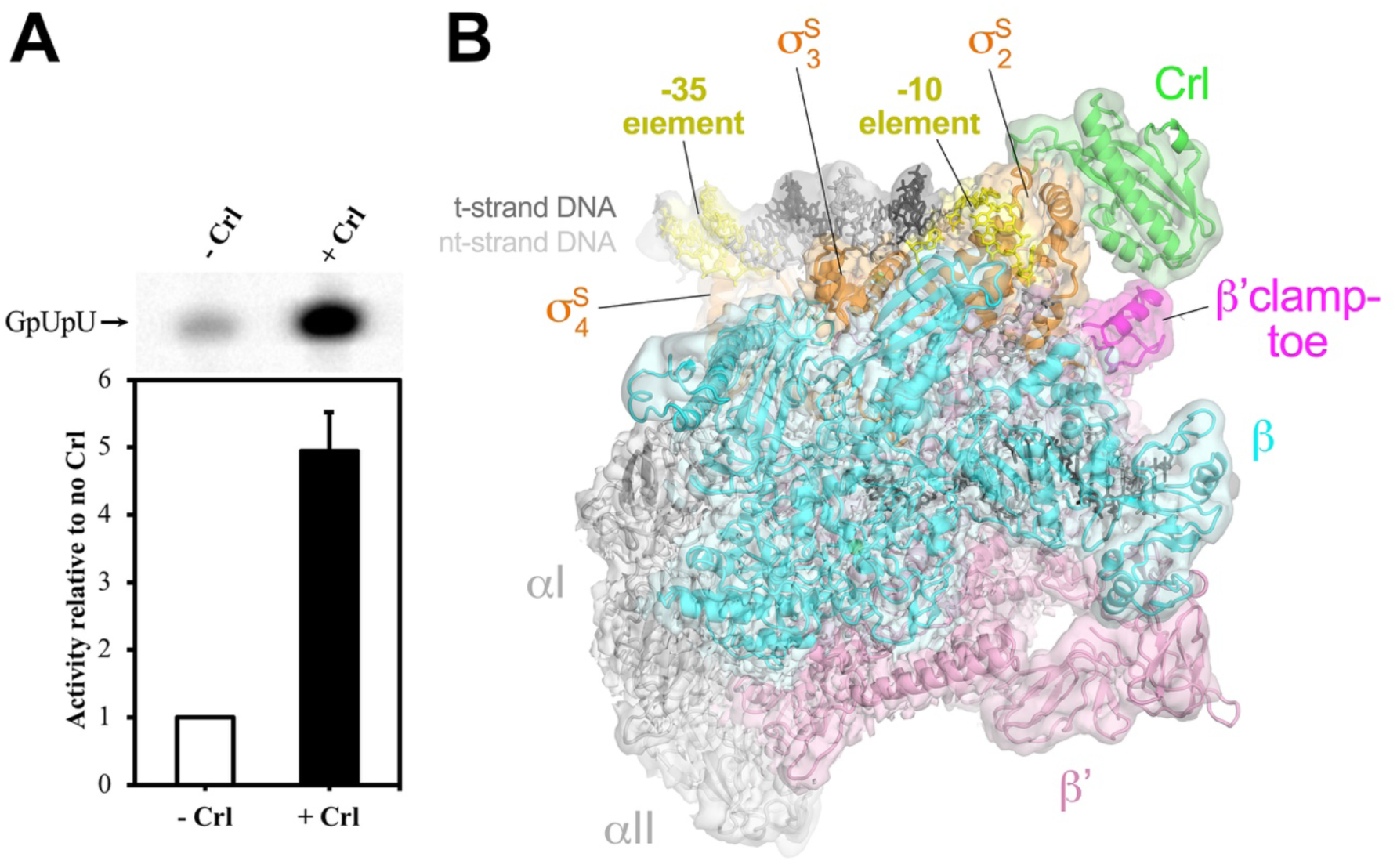
Cryo-EM structure of Crl-Eσ^S^-*dps*-RPo. (**A**) *Sty* Crl with *Sty* σ^S^ activates transcription by *Eco* RNAP on the *Eco dps* promoter. *top. In vitro* abortive initiation of RNA trinucleotide synthesis using a *dps* promoter template. Transcription was initiated with GpU RNA dinucleotide and α-^32^P-UTP. The radioactive GpUpU product was visualized by denaturing polyacrylamide gel electrophoresis and autoradiography. *bottom*. Plotted is RNA trinucleotide synthesis with Crl (+Crl) relative to no Crl (-Crl). The error bars denote standard deviation of three measurements. (**C**) The 3.3 Å nominal resolution cryo-EM density map of Crl-Eσ^S^-*dps*-RPo is rendered as a transparent surface and colored as labeled. The map is low-pass filtered according to the local resolution (53). Superimposed is the final refined model. The proteins are shown as backbone ribbons, and the nucleic acids are shown in stick format.

### Cryo-EM structure of Crl-Eσ^S^-*dps*-Rpo

To prepare a complex for structure determination, we incubated Crl-Eσ^S^ with a duplex *dps* promoter construct (−46 to +20) containing a non-complementary ‘seed’ bubble from −7 to −4 (*SI Appendix*, Fig. S1A) to pre-nucleate the transcription bubble and favor the formation of a homogenous open promoter complex (RPo) as described previously (33). The entire 477 kDa complex (Crl-Eσ^S^-*dps*-RPo) was purified by size-exclusion chromatography (*SI Appendix*, Figs. S1B, C) and cryo-EM grids were prepared as described in Materials and Methods.

The structure of the complex was determined by single-particle cryo-EM (Fig. 1B). Analysis of the cryo-EM data yielded a single structural class (*SI Appendix*, Fig. S2) at a nominal resolution of 3.3 Å, ranging from 2.8 Å in the well-ordered core of the complex to 6.5 Å at the flexible periphery (*SI Appendix*, Fig. S3). A structural model was built and refined into the cryo-EM map (Fig. 1B; *SI Appendix*, Table S2). Initial examination of the cryo-EM structure revealed three key features (Fig. 1B): First, expected interactions occurred between σ^S^_4_ and the −35 promoter element, which were not observed in a previously determined crystal structure of a σ^S^ transcription initiation complex due to crystal packing restraints (PDB 5IPL) (33). Second, Crl bound σ^S^_2_ in a manner predicted from the results of previous studies (26, 38) and is located at the periphery of the complex near the upstream edge of the transcription bubble. Third, Crl also interacted with the RNAP β’ subunit.

### Structural basis for selective activation of σ^S^ by Crl

The σ^S^ is the closest relative of σ^70^ in terms of sequence, domain architecture, and promoter recognition properties (39). Our structure and the Eσ^S^-RPo crystal structure (33) revealed the expected structural similarity between domains 2, 3, and 4 of σ^S^ and σ^70^ (*SI Appendix*, Fig. S4). Despite these similarities, Crl specifically activates σ^S^. The main difference between σ^S^ and σ^70^ is the non-conserved region (NCR) of σ^70^, a 250 amino acid insertion located between regions 1.2 and 2.1 that is absent in σ^S^ (*SI Appendix*, Figs. S4, S5).

As previously reported, Crl is small arc-shaped protein with a shallow concave surface composed of four antiparallel β-strands and flanked by intervening loops (34, 35). This cavity makes extensive electrostatic, polar, and hydrophobic interactions with helix α2 (A73 to R85) of σ^S^, which resides within conserved region σ^S^_1.2_ (3) (*SI Appendix*, Fig. S4). In σ^70^, the corresponding helix extends to become part of the NCR and would sterically clash with Crl binding (26), explaining why Crl selectively binds and activates σ^S^ but not σ^70^.

The central role of σ^S^ helix α2 confirms previous reports that identified key residues in σ^S^ for its interaction with Crl. *Sty* σ^S^ R82 is not conserved in *Pseudomonas aeruginosa* (*Pae*) σ^S^, and *Pae* σ^S^ does not interact with *Sty* Crl in *in vivo* bacterial two-hybrid assays unless the corresponding amino acid (Leu) is mutated to an arginine (38). Mutations at this site lead to bacterial colony morphology changes, which is consistent with the interaction of Crl and σ^S^ being important for processes like biofilm formation. In our structure, *Sty* σ^S^ R82 is positioned towards the central cavity of Crl and forms an extensive network of electrostatic, polar, and hydrophobic interactions with Crl-P21, Y22, I23, D36, and C37 (Fig. 2). Our structure also validates the importance of other sites in helix α2 like Y78 and F79, which have been substituted with benzoyl-L-phenylalanine (BPA) and shown to crosslink to Crl (26).

**Fig. 2.**
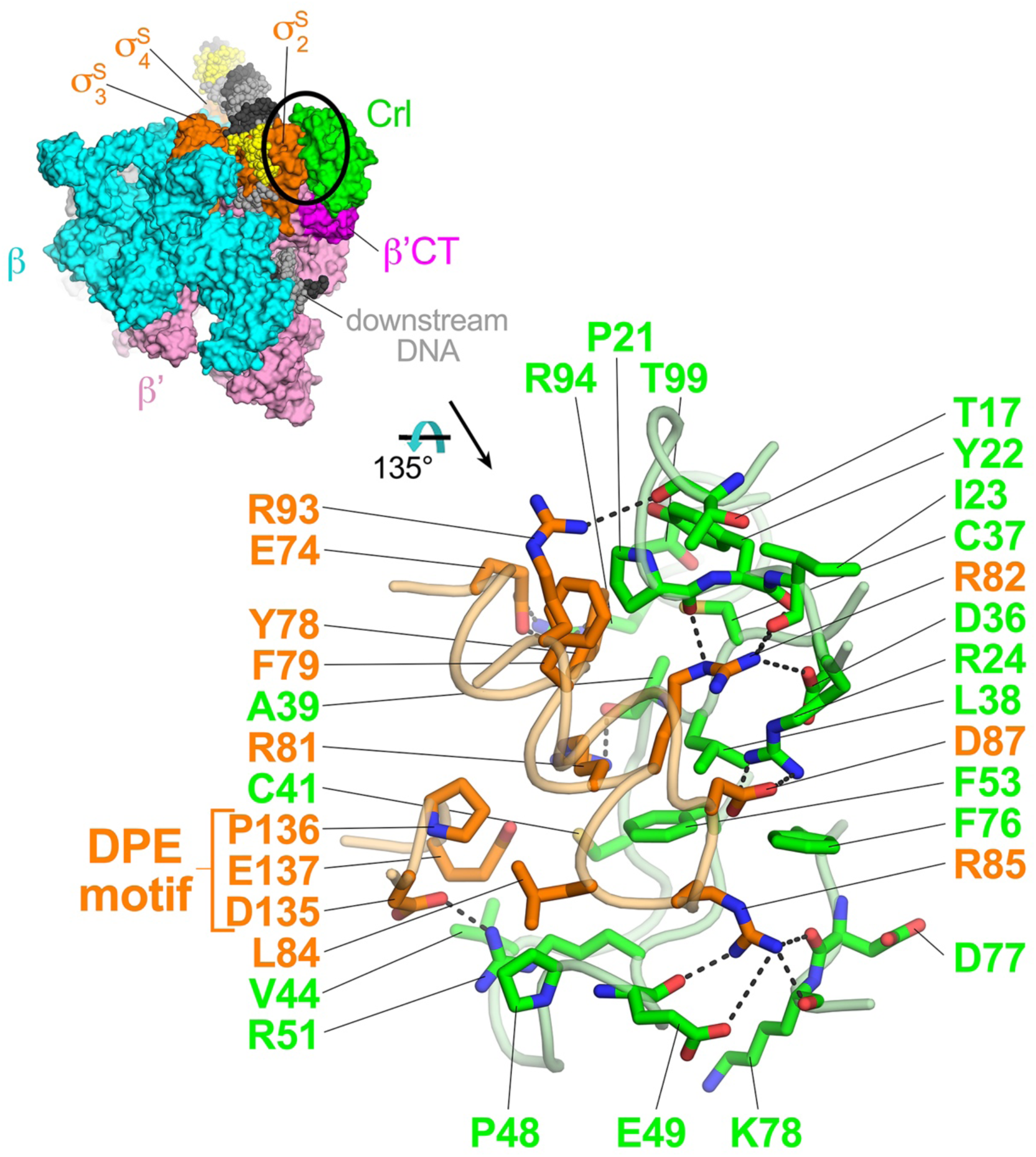
Crl-σ^S^2 interactions. *top left.* The overall structure of Crl-Eσ^S^-*dps*-RPo. Proteins are shown as molecular surfaces with subunits colored as labeled. The DNA is shown as Corey-Pauling-Koltun (CPK) spheres and colored according to Fig. 1B. The circled region is magnified below. *bottom right.* Crl-σ^S^2 interactions. Crl (green) and σ^S^2 (orange) are shown as backbone worms. Residues that interact are shown in stick format. Polar interactions are denoted by dashed gray lines.

Adjacent to σ^S^ helix α2 is a loop within σ^S^_2.3_ that also makes significant interactions with Crl, in particular σ^S^ residues D135 and E137 (Fig. 2). D135 makes favorable electrostatic interactions with Crl R51, which is absolutely conserved among Crl homologs (40). Substitutions Crl R51A or R51K were totally defective in Crl function *in vitro* and *in vivo* (34). D135 and E137 of σ^S^, along with P136, have been referred to as the DPE motif (Fig. 2) and is a key difference from σ^70^ (26). In bacterial two-hybrid assays, substitutions in the DPE motif significantly decrease the interactions between Crl and σ^S^, and a chimeric σ^70^ mutant containing this region of σ^S^ can interact with Crl (26). Thus, substitutions that alter either partner of the Crl/σ^S^-DPE motif interaction interface highlight the importance of this interaction for Crl function.

Altogether, the interaction between Crl and σ^S^ forms an interfacial area of 785 Å^2^ (41) and is completely consistent with previous analyses of Crl-σ^S^ interactions (26, 34, 38, 40). In summary, our structure: 1) confirms the σ^S^ residues previously proposed to interact with Crl based on genetic and biochemical data; 2) identifies additional residues in σ^S^ and Crl involved in the intermolecular interaction; 3) reveals that 6 out of 11 (55%) of the residues of σ^s^ contacting Crl are not conserved in σ^70^ (Fig. 2; *SI Appendix* Fig. S4); and 4) shows that the NCR of σ^70^ would sterically clash with Crl as previously predicted (26) (*SI Appendix*, Fig. S5), thereby explaining the molecular basis of Crl discrimination for regulating σ^S^ and not σ^70^.

### Crl tethers σ^S^ to core RNAP to help activate transcription

In addition to extensive contacts with σ^S^2 (Fig. 2), Crl in the Crl-Eσ^S^*-dps*-RPo structure interacts with a small domain of the *Eco* RNAP β’ subunit we call the β’clamp-toe (β’CT; β’ residues 144-179; Figs. 1B, 3). A sequence alignment of evolutionarily diverse Crl homologs reveals conservation of basic amino acids in the region of Crl that interacts with the β’CT, corresponding to *Sty* Crl K9, R11, K14, and K15 (Fig. 3A). The β’CT is not strictly conserved among bacterial RNAP β’ subunits as it is the site of lineage-specific-insertions in many bacterial clades, including Deinococcus-Thermus and Actinobacteria (42). However, the sequence of the β’CT is conserved among RNAP β’ sequences from *γ*-proteobacteria, including residues that interact directly with Crl: *Eco* β’L166, as well as two acidic residues, D167 and E170 (Fig. 3A). The α-helix of the β’CT that interacts with Crl harbors five conserved acidic residues corresponding to *Eco* β’ E162, E163, D167, E170, and E171 (Fig. 3A). The observation that Crl interacts with core RNAP was verified in photocrosslinking experiments. Crl crosslinked to β’ with BPA substitutions at D167 & F172 but not E142 in a σ^S^-dependent manner, consistent with our structure (Figs. 3B, C).

**Fig. 3.**
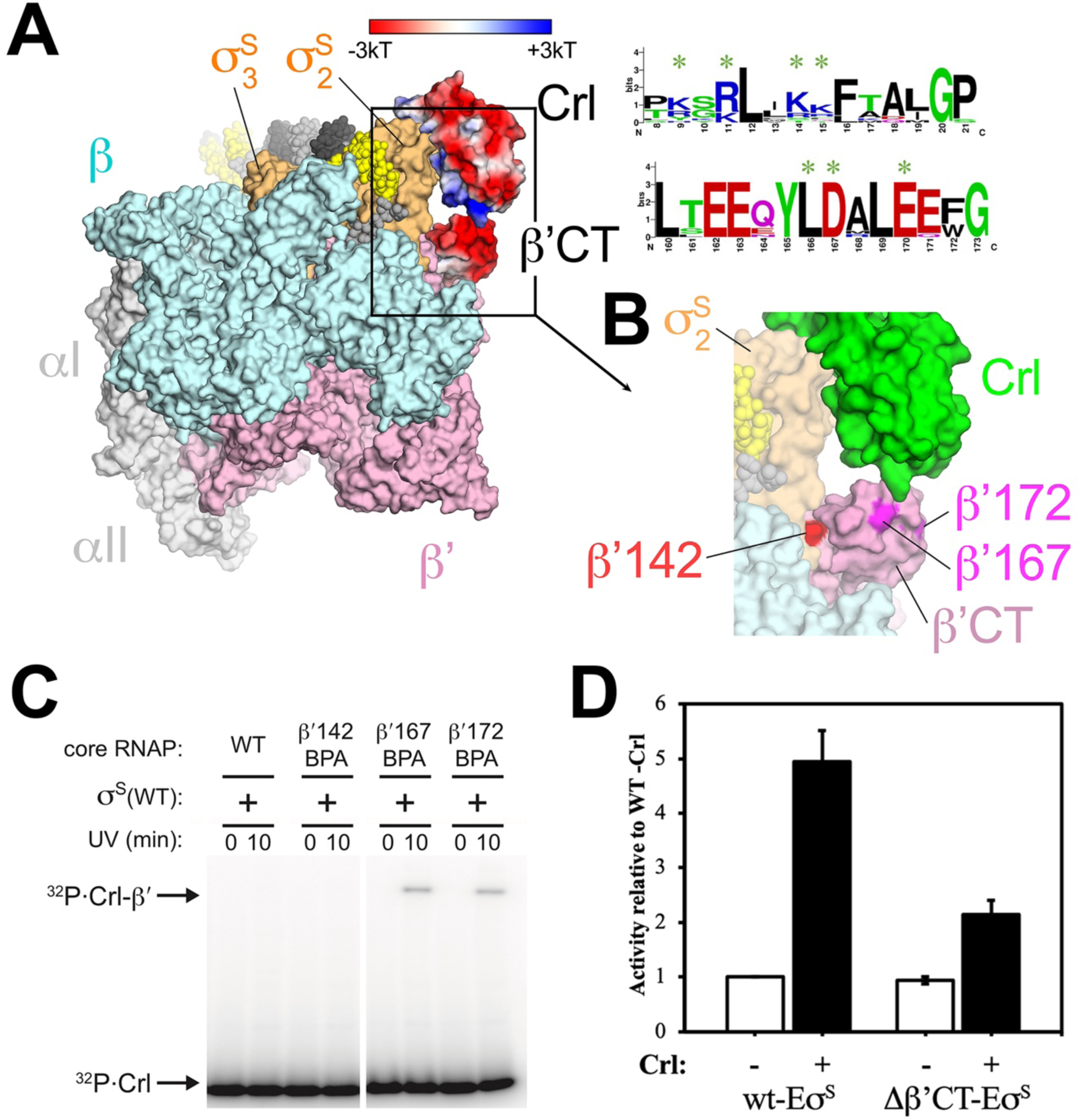
The Crl-β’CT interaction. **(A)** The overall structure of Crl-Eσ^S^-*dps*-RPo. Proteins are shown as molecular surfaces with subunits colored as labeled, except Crl and the β’CT are colored according to electrostatic surface potential (red −3kT; blue, +3kT) (54). The DNA is shown as CPK spheres and colored according to Fig. 1B. On the right are shown sequence logos (55) for the regions of Crl (top) and the β’CT (bottom) that interact with each other. Residues that directly interact are denoted by a green ‘*’ above. The rectangle denotes a region of the structure magnified in (**B**). The sequence logos were derived from sequence alignments of Crl and the RNAP β’ subunit from the same 51 evolutionarily diverse *γ*-proteobacteria (56). **(B)** Magnified view from (**A**) focusing on Crl (green) – β’CT (pink) interaction. **(C)** SDS-PAGE gel showing effect of UV exposure on RNAP core with β’ BPA substitutions incubated with ^32^P-Crl. Crosslinked complexes and free ^32^P-Crl are indicated. RNAP β’ BPA substitutions at residues 167 and 172 crosslink to Crl [magenta in (**B**)]; BPA substitution at 142 [red in (**B**)] does not crosslink to Crl. **(D)** The β’CT is required for full Crl activation. Plotted is RNA trinucleotide synthesis without Crl (-, white bars) or with Crl (+, black bars) for wt-Eσ^S^ or Δβ’CT-Eσ^S^. The values are normalized with respect to wt-Eσ^S^(-Crl). The error bars denote standard deviation of three measurements.

These observations suggest that Crl may assist Eσ^S^ assembly by interacting with σ^S^_2_ and RNAP simultaneously. To test this hypothesis, we investigated Crl function with a mutant RNAP in which the entire β’CT was deleted (Δβ’CT-E) using the quantitative abortive initiation assay with the *dps* promoter. While wt-Eσ^S^ and Δβ’CT-Eσ^S^ had essentially the same transcription activity in the absence of Crl, the presence of a saturating concentration of Crl activated wt-Eσ^S^ ∼5-fold compared to only ∼2.3-fold with Δβ’CT-Eσ^S^ (Fig. 3D). We conclude that the simultaneous interaction of Crl with σ^S^_2_ and the β’CT helps tether σ^S^ to the core RNAP, increasing the stability of Eσ^S^, accounting for partial, but not full, transcription activation function of Crl.

## Discussion

Our results provide insights into the mechanisms by which Crl promotes Eσ^S^ assembly. The Crl-Eσ^S^-*dps*-RPo cryo-EM structure is consistent with, and expands upon, previous information about the interactions of Crl with σ^S^_2_ (26, 28, 34, 35, 40), and includes an interaction with the RNAP β’CT. To our knowledge, this region of core RNAP has not been previously described as a binding determinant for transcription factors and may represent a target for other uncharacterized transcription factors. Deletion of the β’CT had little effect on basal transcription (-Crl) but Crl was unable to fully activate the Δβ’CT-RNAP, suggesting that one mechanism by which Crl activates Eσ^S^ transcription is to stabilize the Eσ^S^ complex by tethering σ^S^ and RNAP. This mechanism is consistent with previous studies showing that Crl activation function is most pronounced when σ^S^ concentrations are low (28, 30).

The tethering mechanism only accounts for partial Crl activity (Fig. 3D). Our studies do not exclude a post-Eσ^S^ assembly role for Crl in transcription activation, such as facilitating promoter melting (RPo formation) or promoter escape. Crl was shown to promote full transcription bubble formation at the *Sty katN* promoter, particularly at 20°C where Eσ^S^ without Crl could only form partially melted intermediates (43). In our structure, Crl does not interact with the promoter DNA. However, the σ^S^ DPE motif (D135/P136/E137), critical for the Crl-σ^S^ interaction (Fig. 2), is part of the conserved region 2.3 of σ^S^, comprising a short loop that forms a part of the binding pocket for the nontemplate strand -11A, the most conserved position of bacterial promoters (44, 45). In fact, the -11A base forms a hydrogen-bond with the α-carbon backbone NH of σ^S^ D135, the side chain of which interacts with Crl (Fig. 4). Thus, Crl might stabilize a conformation of the σ^S^ DPE motif that facilitates transcription bubble formation.

**Fig. 4.**
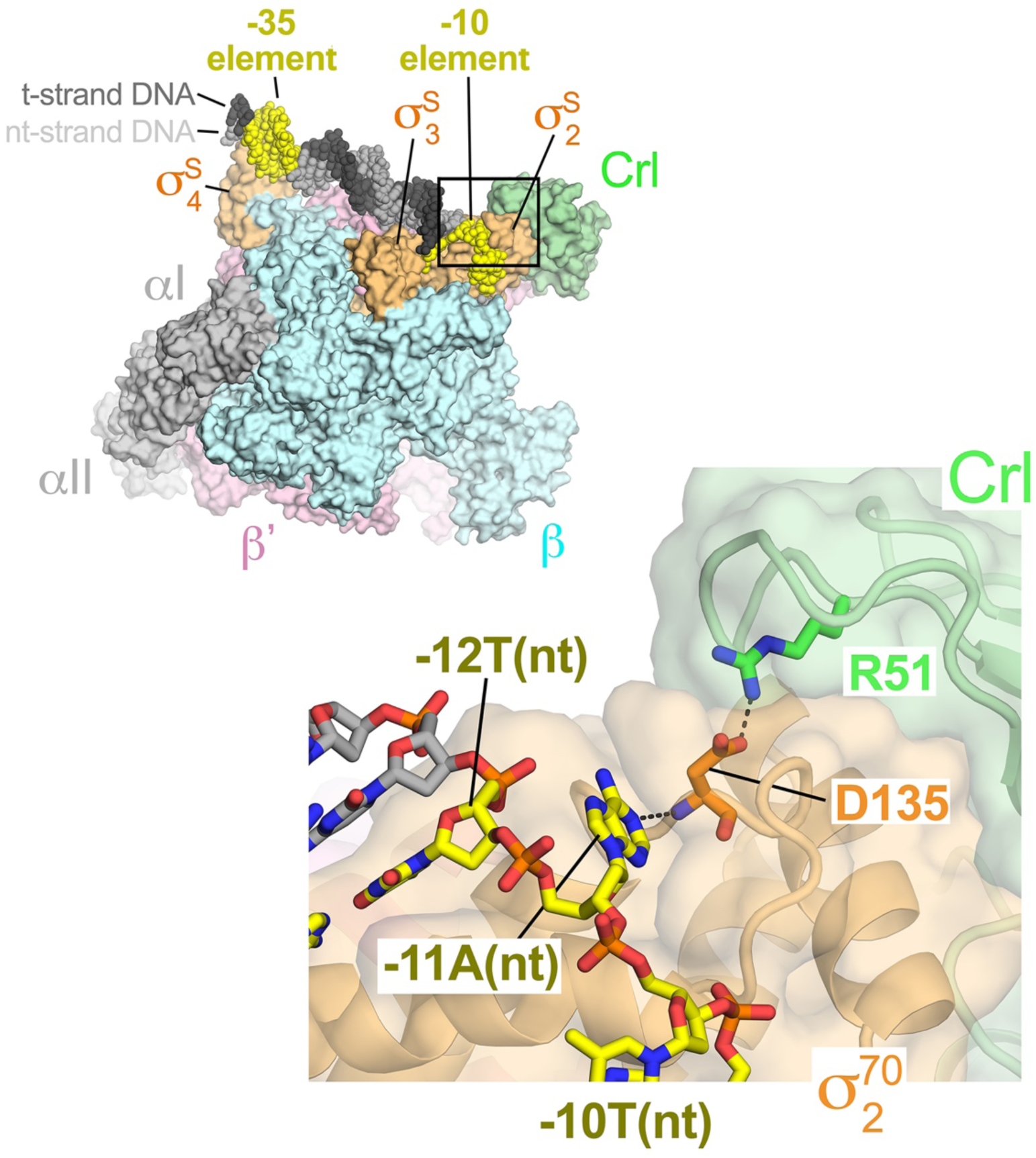
Crl interacts with σ^S^ residues involved in non-template strand -11A capture. *top left.* The overall structure of Crl-Eσ^S^-*dps*-RPo. Proteins are shown as molecular surfaces with subunits colored as labeled. The DNA is shown as CPK spheres and colored according to Fig. 1B. The boxed region is magnified below. *bottom right.* Crl (light green) and σ^S^_2_ (light orange) are shown as backbone worms. The promoter DNA is shown in stick format. The side chains of Crl-R51 (green) and σ^S^-D135 (orange) are also shown in stick format. The side chain and backbone amide of σ^S^-D135 make simultaneous polar interactions with Crl-R51 and -11A(nt), respectively (denoted by dashed gray lines).

Our results suggest Crl exerts direct transcription activation through contacts with the β’CT. The β’CT was previously shown to interact with the σ^70^NCR, which antagonized the σ^70^-β’clamp-helices interaction to enhance promoter escape and reduce early elongation pausing (46). The roles of Crl in promoter escape and early elongation pausing have not been examined, but the similarity in spatial arrangement between Crl and the σ^70^_NCR_ with respect to the β’CT point towards possible analogous roles. Furthermore, the conservation of the β’CT in Crl-containing *γ*-proteobacteria (Fig. 3A) highlights its mechanistic importance and suggests it might be the target of regulation by other transcription factors or play other important roles.

Previous studies determined crystal structures of Eσ^S^ bound to a promoter fragment (highest resolution of 3.6 Å; 5IPL) containing σ^70^ consensus −10 and −35 elements (33). These structures showed the engagement of Eσ^S^ with the promoter −10 element and downstream part of the promoter, but the upstream promoter −35 element was not bound to σ^S^_4_due to exclusion by crystal packing. The protein components of our structure align well with this previous structure (0.96 rmsd over 2,600 Cα’s), and σ^S^_2_ (σ^S^ residues 53-167) aligns with an rmsd of 0.51 Å over 104 Cα’s, indicating that Crl binding does not induce significant conformational changes in σ^S^_2_ once Eσ^S^ is formed. Similarly, Crl in the Crl-Eσ^S^-*dps*-RPo cryo-EM structure aligns with an rmsd of 0.8 Å over 113 Cα’s with the Crl crystal structure [3RPJ; (34)].

Our cryo-EM structure contains a *dps* promoter DNA fragment. The *dps* promoter is part of the σ^S^ regulon (36, 47). The *dps* promoter is transcribed by both Eσ^70^ and Eσ^S^ *in vitro* but shows a marked preference for Eσ^S^ (37). However, our RPo structure does not reveal striking differences in promoter DNA interactions comparing σ^70^ and σ^S^, which suggests that differences in promoter preference may not be simply due to σ-promoter interactions in the final RPo, but to differences in the kinetics of RPo formation. Since Crl is induced at low temperatures, it would be interesting to determine if activation by Crl has more mechanistic functional importance for transcription initiation at low temperature where promoter melting by RNAP is inhibited (27, 43).

Transcription activators Crl, GcrA, GrgA, and RbpA interact with domain 2 and/or the NCR of σ factors and represent an emerging paradigm in bacterial transcription regulation (48-51). Crl plays a unique role as it specifically activates transcription of σ^S^-dependent genes, which bacteria express in order to respond to changes in their environment. We propose that Crl be termed a σ-activator, which may represent a class of factors with similar function. We note that mycobacterial RbpA has been shown to tether σ^A^ to core RNAP via the β’ Zinc binding domain (52). It remains to be seen whether GcrA or GrgA interact with RNAP to tether the σ factors they regulate.

## Materials and Methods

Detailed descriptions of the purification of *Sty* Crl, *Sty* σ^S^, *Eco* ΔαCTD-RNAP, construction and purification of the *Eco* Δβ’CT-RNAP, transcription assays, preparation of Crl-σ^S^-*dps*-RPo for cryo-EM, cryo-EM grid preparation, cryo-EM data acquisition and processing, model building and refinement, and benzoyl-L-phenylalanine crosslinking, are provided in *SI Appendix*.

## Acknowledgements

We thank James Chen and Jin Young Kang, Deena Oren, Tom Walz, and Chris Lima for helpful advice, Yuhong Zuo and Ann Hochschild for sharing material, and M. Ebrahim and J. Sotiris at The Rockefeller University Cryo-EM Resource Center for help with data collection. J. Winkelman constructed some of the RNAP-BPA derivatives used for the photocrosslinking experiments, and P.D. Olinares and B.T. Chait performed native mass spectrometry measurements to confirm the Δβ’CT-RNAP construct. A.J.C. was supported by a Robert D. Watkins Graduate Research Fellowship from the American Society for Microbiology. This work was supported by NIH grants R01 GM347048 to R.L.G., R01 GM114450 to E.A.C., and R35 GM118130 to S.A.D.

