## Supplemental Information for "Structural basis for transcription activation by Crl through tethering of σ^S^ and RNA polymerase"

### Supplementary Information

#### Expanded Materials and Methods

##### Protein purification

***Sty* Crl.** $His_6$-ppx-Crl was recombinantly expressed from a pET-21a plasmid transformed into *Eco* BL21 LOBSTR strain, which removes common contaminants in His-tag purifications (1). LB media (2-liters) was inoculated with cell colonies grown overnight on LB agar plates with 100 μg/ml. Cultures were incubated with shaking at 200 rpm at 37$^{o}$C until $OD_{600nm}$ = 0.4 when the temperature was reduced to 30$^{o}$C and at $OD_{600nm}$ = 0.6 protein expression was induced by adding isopropyl β-D-1-thiogalactopyranoside (IPTG, final concentration 0.5 mM) with further incubation at 30°C for 2 hours. Cells were harvested and resuspended in 60 mL of 200 mM NaCl, 50 mM Tris-HCl, pH 8.0, 10 mM imidazole, 5% glycerol (v/v). Cells were lysed using a continuous flow French press (Avestin) at 15,000 psi. Cell lysate was centrifuged at 15000 rpm (~12000 g) for 40 minutes. Cell lysis supernatant was loaded into a 5 mL $Ni^{2+}$-charged HiTrap IMAC column (GE Healthcare). The column was washed with 200 mL of 30 mM imidazole, then proteins were eluted with 300 mM imidazole. PreScission protease (GE Healthcare) was added and the sample was dialyzed (8 kDa molecular weight cutoff dialysis tubing) against 1 L of 200 mM NaCl, 50 mM Tris-HCl, pH 8.0, 5% glycerol (v/v). After overnight incubation, a subtractive IMAC was performed to separate untagged Crl from the cleaved $His_6$-tag and uncleaved $His_6$-Crl. The untagged Crl was further purified by size exclusion chromatography (SEC) on a Superdex 200 HiLoad column (GE Healthcare). Fractions containing Crl were pooled, concentrated to 160 μM by centrifugal filtration (Amicon Ultra), supplemented with glycerol to 20% (v/v), then divided into 40 μL aliquots and stored at -80°C.

***Sty* $\sigma^S$.** Gibson assembly (2) was used to generate $His_{10}$-SUMO-$\sigma^S$ in pET-21a. The Crl purification protocol was used for this construct, except the $His_{10}$-SUMO-tag was removed using ULP-1. Protein was divided into 40 μL of 200 μM aliquots in the presence of 20% glycerol and stored at -80°C.

***Eco* ΔαC-terminal domain RNAP: Eco** RNAP lacking the α C-terminal domains was expressed and purified as previously described (3).

**Construction of RNAP β' ΔCT mutant**. We generated an *Eco* RNAP expression construct with the β'CT deleted and replaced with a Gly-Ser linker [Δβ'(Tyr144-Lys179)::(Gly144-Ser145)]. The desired deletion was constructed in three steps. First, a plasmid template that encoded the last 57 amino acids in the sequence of the β and the first 625 amino acids of β' was PCR amplified using a primer in the reverse direction before β' CTH, and a forward primer after the β' CTH. Second, the resulting linear PCR product was re-circularized using Gibson assembly (2). Third, this product was cut with SbfI and BsmI and appended into a full RNAP expression plasmid, which encodes the rest of the core RNAP subunits. This expression plasmid is different than the one used for the cryo-EM studies and leads to the expression of RNAP harboring full-length α-subunit. The plasmid encodes full-length wt-α, β, β'(Δβ'CT)-ppx-$His_{10}$, and ω. DNA sequencing and native mass spectrometry confirmed the correct construct was designed and purified. For transcription assays this construct was purified alongside its wt-RNAP parent with full-length β'.

**Transcription assays**. Abortive initiation assays were adapted from previous protocols (4). The reactions were performed at 37°C in a buffer containing 50 mM Tris-HCl, pH 8, 100 mM K-glutamate, 10 mM $MgCl_2$, 10 mM Dithiothreitol (DTT), and 100 μg/mL BSA. Reactions contained final concentrations of 50 nM core RNAP, 100 nM $\sigma^S$, 3.2 μM Crl, and 10 nM *dps* promoter DNA (-39 to +42; Fig. S1A). First, $\sigma^S$ was incubated with Crl or buffer for 10 minutes at 37°C. Next, $\sigma^S$ or Crl-$\sigma^S$ were incubated with wt-RNAP or Δβ'CT-RNAP for 10 minutes at 37°C to form holoenzyme. Lastly, promoter DNA was added to the holoenzymes and incubated for 10 minutes at 37°C to allow RPo formation. Abortive transcription was initiated by adding a mix of GpU dinucleotide primer (250 μM, TriLink), [α-$^{32}$P]UTP (1.25 μCi), and unlabeled GTP (50 μM). After 20 minutes, reactions were quenched with 2X stop buffer (8 M urea, 89 mM Tris-HCl, 89 mM boric acid, 2 mM EDTA, 0.05% bromophenol blue, 0.05% xylene cyanol) and separated on a 23% (w/v) urea-polyacrylamide gel. Abortive products were visualized by phosphorimaging and quantified using ImageJ (5).

**Preparation of Crl- $\sigma^S$-*dps*-RPo for Cryo-EM**. A 400 uL sample was prepared containing final concentrations of 40 μM RNAP, 80 μM $\sigma^S$, 160 μM Crl, and 80 μM promoter DNA (Fig. 1B). First, $\sigma^S$ was incubated with Crl for 10 minutes at 37°C. Next, RNAP was added and incubated for 10 minutes at 37°C to form Crl-E$\sigma^S$. Lastly, promoter DNA was added and incubated for 10 minutes at 37°C to allow RPo formation. The sample was injected into a 24 mL

Superose 6 Increase column (GE Healthcare Life Sciences, Pittsburgh, PA) pre-equilibrated with 20 mM Tris-HCl, pH 8.0, 150 mM K-Glutamate, 10 mM $MgCl_2$, 5 mM DTT. Fractions containing the complex were pooled and concentrated to 5 mg/mL protein by centrifugal filtration (EMD-Millipore, Darmstadt, Germany). An RNA oligonucleotide (CUCG) was added to a final concentration of 20 μM. CHAPSO (3-([3-cholamidopropyl]dimethylammonio)-2-hydroxy-1-propanesulfonate) was then added to a final concentration of 8 mM (6) and the sample was kept at room temperature before grid preparation.

**Cryo-EM grid preparation**. C-flat holey carbon grids (CF-1.2/1.3-4Au) were glow-discharged for 30 s before the application of 3.5 μl of the sample. After 4 seconds the grids were plunge-frozen in liquid ethane using an FEI Vitrobot Mark IV (FEI, Hillsboro) with 100% chamber humidity at 25°C. Grids were stored in liquid nitrogen.

**Cryo-EM data acquisition and processing**. Grids were imaged using a 300 keV Titan Krios (FEI) microscope equipped with a K2 Summit direct electron detector. Images were collected using Serial EM (7) in super-resolution counting mode with a super-resolution pixel size of 0.65 Å (22,500 X) and a defocus range of 0.8 to 2.4 μm with 0.2 μm steps. Images were collected at a dose rate of 8 electron/pixel/s. 50 frames were collected over 15 seconds (0.3 s/frame), yielding a dose of 71 electron/Å$^2$. Dose-fractionated frames were 2 X 2 binned (giving a pixel size of 1.3 Å).

Data processing was initially carried out in Relion3 using MortionCor2 to bin and align the 50 frames (8, 9) to obtain single dose filtered micrographs. Particles in the micrographs were automatically picked using Gautomatch (K. Zhang, MRC Laboratory of Molecular Biology, Cambridge, UK, http://www.mrc-lmb.cam.ac.uk/kzhang/Gautomatch), which resulted in ~658,000 particles. We built an initial model using a previously published structural model of a $\sigma^S$-TIC [PDB 5IPL; (10)] with a docked model of Crl from an x-ray crystal structure [PDB 3RPJ; (11)]. The particles were classified into three classes using this model. One class was refined to 4.6 Å, the particles from the other two classes were discarded. The promising class was furthered classified into 3 additional classes. Two of the classes showed good reconstructions and they were combined to obtain ~292,000 high quality, while particles from the third class were discarded. A series of five iterative 3D auto-refinements of the high-quality particles were conducted using particle CTF refinement and Bayesian particle polishing in Relion3. The final polished particles were imported into CryoSPARC2 (12) to perform a homogenous refinement, followed by a local refinement, which yielded the final density map with an overall resolution of 3.26 Å. Local resolution calculations were also conducted using cryoSPARC.

**Model building and refinement**. An initial structural model was assembled from the core RNAP component of PDB 5VT0 (13), *Eco* $\sigma^S$ from PDB 5IPL (10), *Proteus mirabilis* Crl [PDB 3RPJ; (11)], and DNA from PDB 4XLN (14). The $\sigma^S$, Crl, and DNA components were mutated to correspond to *Sty* $\sigma^S$, *Sty* Crl, and the *dps* promoter construct used for cryo-EM (Fig. 1B). The model was fit into the cryo-EM density map using UCSF Chimera (15). Appropriate domains of the complex were rigid-body refined, then subsequently refined with secondary structure and nucleic acid restraints using PHENIX real space refinement (16), along with rounds of manual adjustment using COOT (17).

**Benzoyl-L-phenylalanine crosslinking.** Benzoyl-L-phenylalanine (BPA)-mediated crosslinking was carried out using modifications of a previously described procedure (11). *Eco* $\sigma^S$ and *Eco* N-HMK-Crl were purified and HMK-Crl was $^{32}P$-labeled as described (11). WT or BPA-substituted *Eco* core RNAPs were overexpressed in BL21DE3 from multisubunit plasmid pIA299 derivatives and purified as described (18) (pRLG7814, WT RNAP; pRLG11773, β'142BPA RNAP; RLG11777, β'167BPA RNAP, pRLG13513, β'172BPA RNAP). $\sigma^S$ and Crl were preincubated for 10 min in buffer containing 40 mM Tris-HCl, pH 8, 100 mM KCl, 10 mM $MgCl_2$, 5 mM DTT and 100 µg/ml BSA. RNAP core enzyme was then added and incubated for an additional 10 min to assemble $E\sigma^S$. Samples were exposed to UV (360 nm) for 0-10 min (as indicated), mixed with 4 µl 4X LDS (Invitrogen) + 25 mM DTT, heated at 85°C for 3 min, loaded onto a 4-12% Bis-Tris gel (Invitrogen) and run with 1X MES buffer at 120V for 1.5-2 hrs. Gels were transferred to Whatman paper and dried for 1 h at 80°C under vacuum. Crosslinked proteins were visualized using a Typhoon 9400 and ImageQuant software (GE Healthcare).

#### Supplementary Tables

**Table S1. Sequence identity between *Eco* and *Sty*.**

| Subunit | Alignment length (residues) | % sequence Identity |
|---|---|---|
| α | 329 | 100.0 |
| β | 1342 | 98.7 |
| β' | 1407 | 98.6 |
| ω | 91 | 100.0 |
| $\sigma^{70}$ | 615 | 97.6 |
| $\sigma^{S}$ | 330 | 99.1 |
| Crl | 133 | 84.2 |
| overall | 4,576 | 98.3 |

**Table S2. Cryo-EM data collection, refinement and validation statistics**

| | Crl-E$\sigma^S$ RPo (EMDB-20090) (PDB 6OMF) |
|---|---|
| **Data collection and processing** | |
| Magnification | 22,500 |
| Voltage (kV) | 300 |
| Electron exposure (e–/Å$^2$) | 71 |
| Defocus range (μm) | 0.8 – 2.4 |
| Pixel size (Å) | 1.3 |
| Symmetry imposed | C1 |
| Initial particle images (no.) | 656,075 |
| Final particle images (no.) | 292,588 |
| Map resolution (Å) - FSC threshold 0.143 | 3.3 |
| Map resolution range (Å) | 2.8 - 6.5 |
| **Refinement** | |
| Initial model used (PDB code) | RPo, 5IPL/Crl, 3RPJ |
| Model resolution (Å) FSC threshold 0.5 | 5IPL 3.6, Crl 1.9 |
| Model resolution range (Å) | 3.1 – 7 |
| Map sharpening *B* factor (Å$^2$) | 141.1 |
| Model composition | |
| Non-hydrogen atoms | 30,250 |
| Protein residues | 3,616 |
| Nucleic acid residues | 92 |
| Ligands | 3 (1 $Mg^{2+}$, 2 $Zn^{2+}$) |
| *B* factors (Å$^2$) | |
| Protein | 144.3 |
| Nucleic acid | 219.7 |
| Ligands | 143.1 |
| R.m.s. deviations | |
| Bond lengths (Å) | 0.011 |
| Bond angles (°) | 0.978 |
| Validation | |
| MolProbity score | 2.78 |
| Clashscore | 9.32 |
| Poor rotamers (%) | 12.31 |
| Ramachandran plot[a] | |
| Favored (%) | 82.5 |
| Allowed (%) | 17.6 |
| Disallowed (%) | 0 |

[a] Ramachandran plot parameters from PROCHECK(19)

#### Supplementary Figures

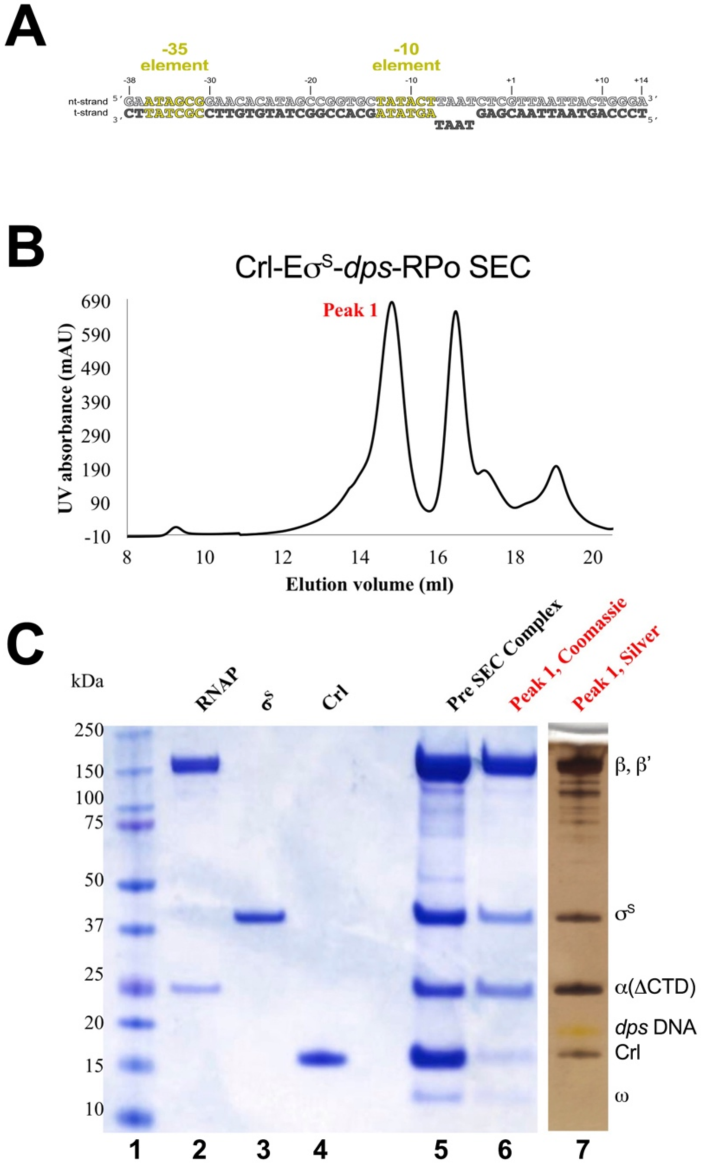

Cartagena et al., Figure S1

**Fig. S1.** Purification of Crl-Eσ$^S$-*dps*-RPo.

(**A**) The *Eco dps* promoter construct used for cryo-EM.

(**B**) Size exclusion chromatography (SEC) profile of Crl-Eσ$^S$-*dps*-RPo. Peak 1 is analyzed by SDS-PAGE in (**B**).

(**C**) SDS-PAGE analysis of Crl-Eσ$^S$-*dps*-RPo. Lane 1, molecular weight markers; lane 2, purified *Eco* core ΔαCTD-RNAP; lane 3, purified *Sty* σ$^S$; lane 4, purified *Sty* Crl; lane 5, SEC load; lane 6, Peak 1 from SEC elution [see (**A**)], Comassie blue stained; lane 7, Peak 1 from SEC elution, silver stained.

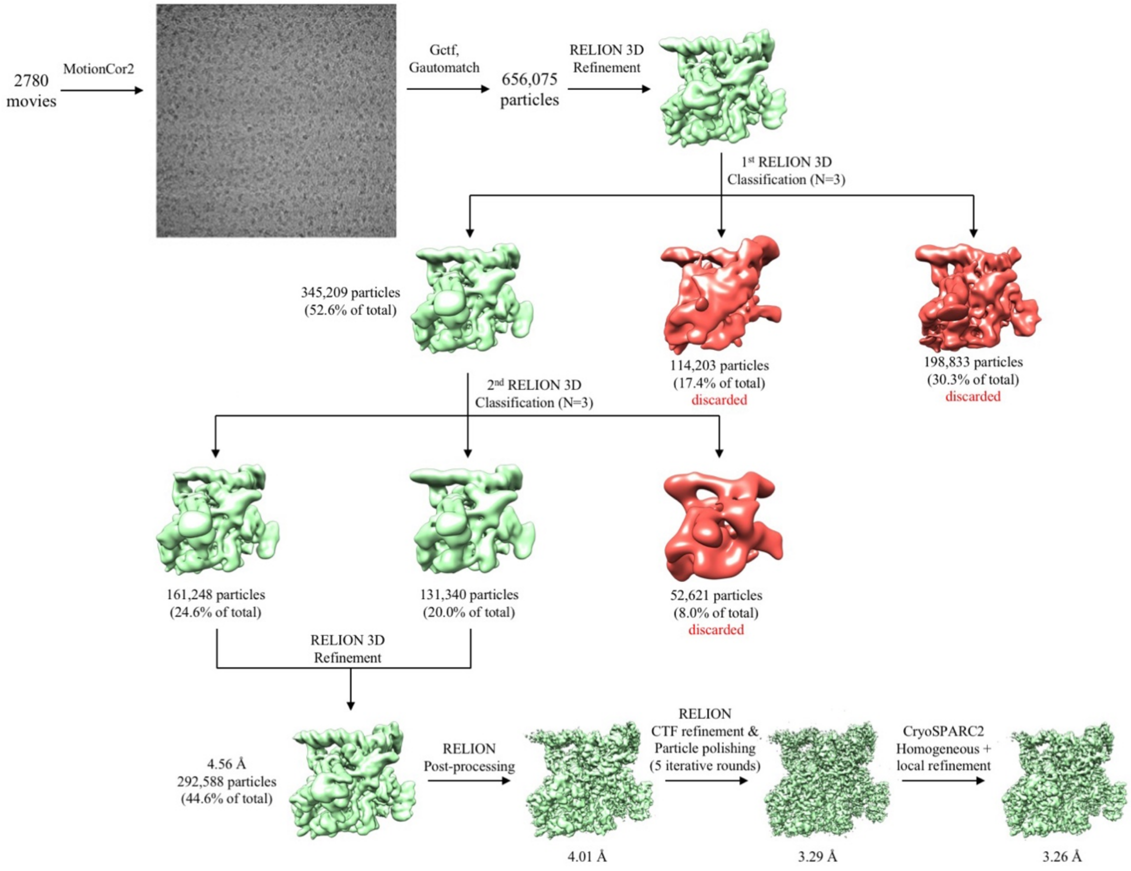

Cartagena et al., Figure S2

**Fig S2.** Crl-Eσ$^{S}$-*dps*-RPo cryo-EM pipeline.

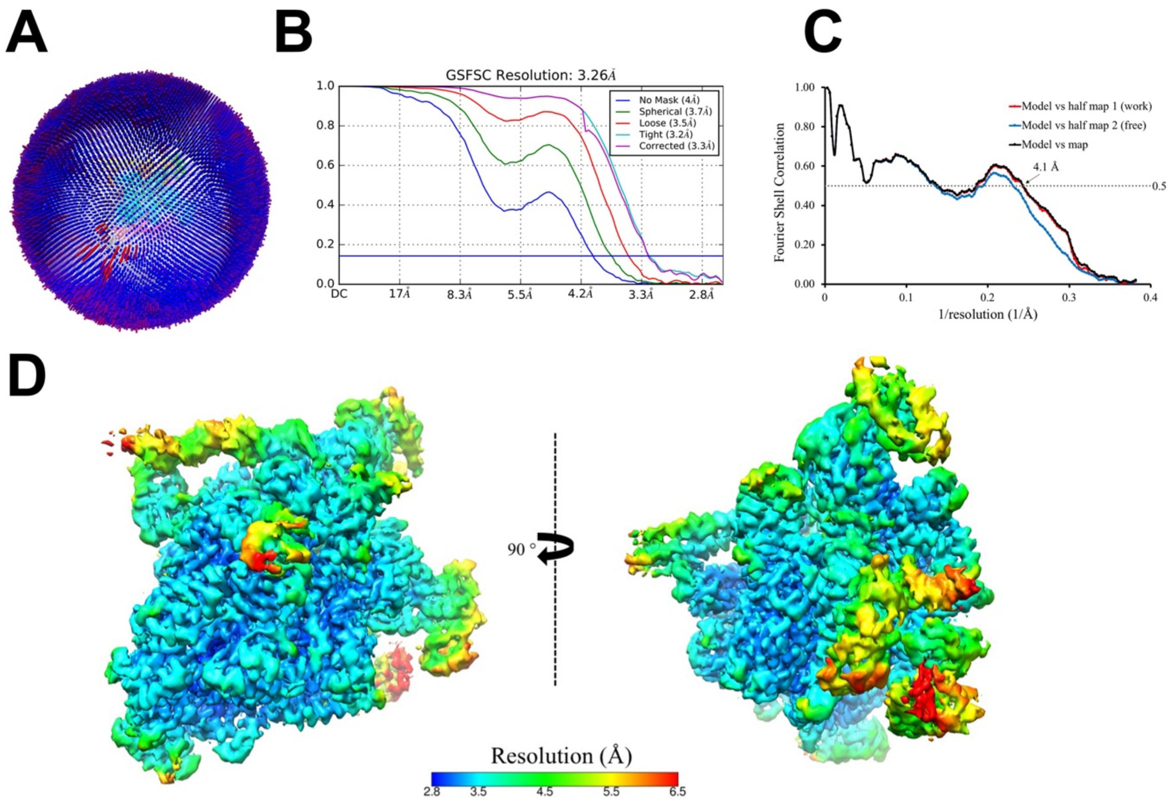

Cartagena et al., Figure S3

**Fig. S3.** Cryo-EM of Crl-Eσ$^{S}$-*dps*-RPo.

(**A**) Angular distribution for Crl-Eσ$^{S}$-*dps*-RPo particle projections.

(**B**) Gold-standard FSC of Crl-Eσ$^{S}$-*dps*-RPo. The gold-standard FSC was calculated by comparing the two independently determined half-maps from Cryosparc. The dotted line represents the 0.143 FSC cutoff, which indicates a nominal resolution of 3.3 Å.

(**C**) FSC calculated between the refined structure and the half map used for refinement (work), the other half map (free), and the full map.

(**D**) The cryo-EM density map of Crl-Eσ$^{S}$-*dps*-RPo is colored according to the local resolution (20).

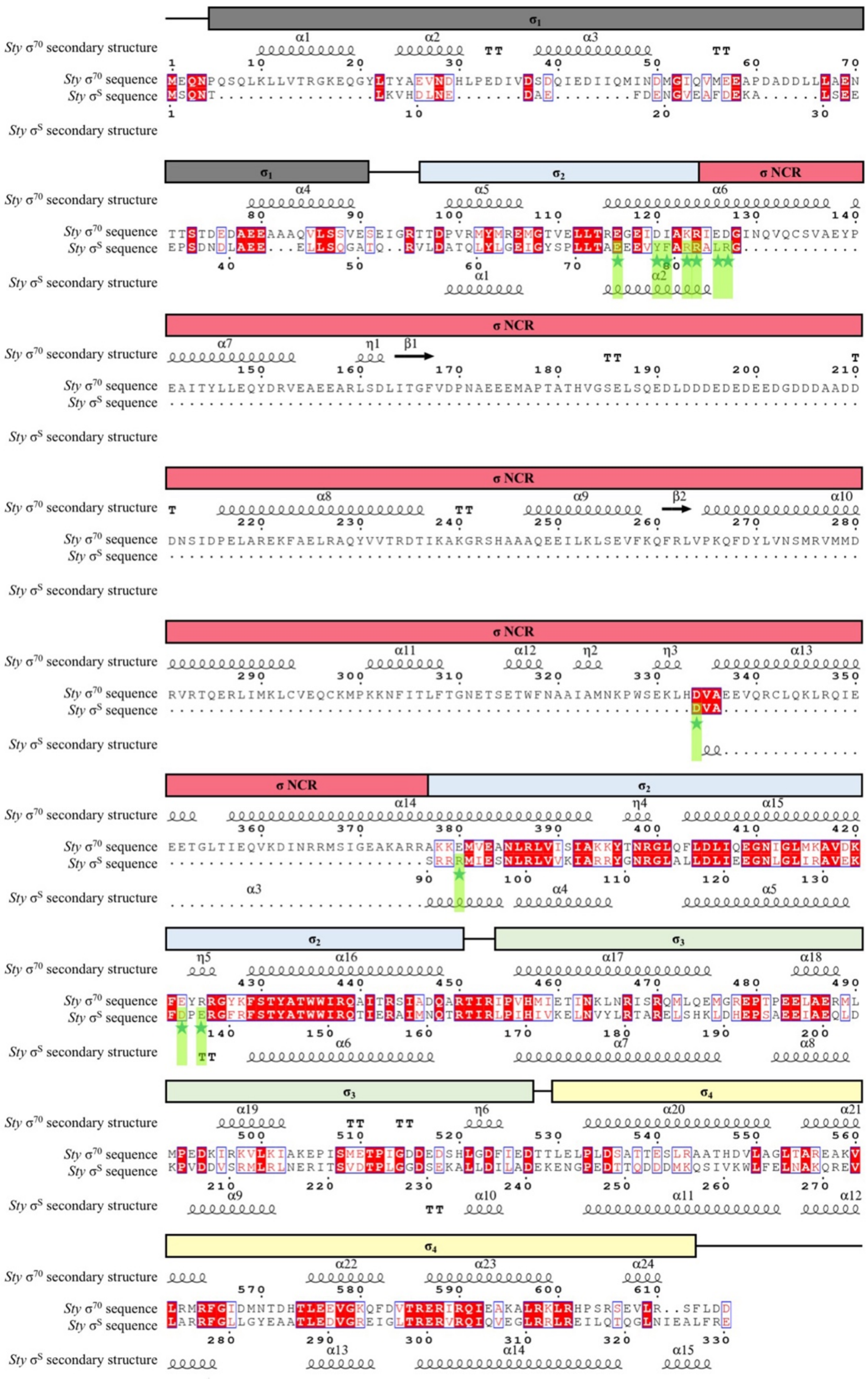

Cartagena et al., Figure S4

**Fig. S4.** *Sty* $\sigma^{70}$ and $\sigma^S$ sequence alignment, secondary, and domain architecture.

Identical residues between the two sequences are boxed red, homologous residues colored red and boxed in white. Above and below the sequences is schematically illustrated the $\sigma^{70}$ (above) *Sty* $\sigma^S$ (below) secondary structures ($\alpha$-helices indicated by coils). Above the secondary structure is illustrated the domain architecture. $\sigma_1$ and $\sigma_{NCR}$ are unique to $\sigma^{70}$. The $\sigma^S$ residues that interact directly with Crl in the cryo-EM structure are marked with green stars and shaded green.

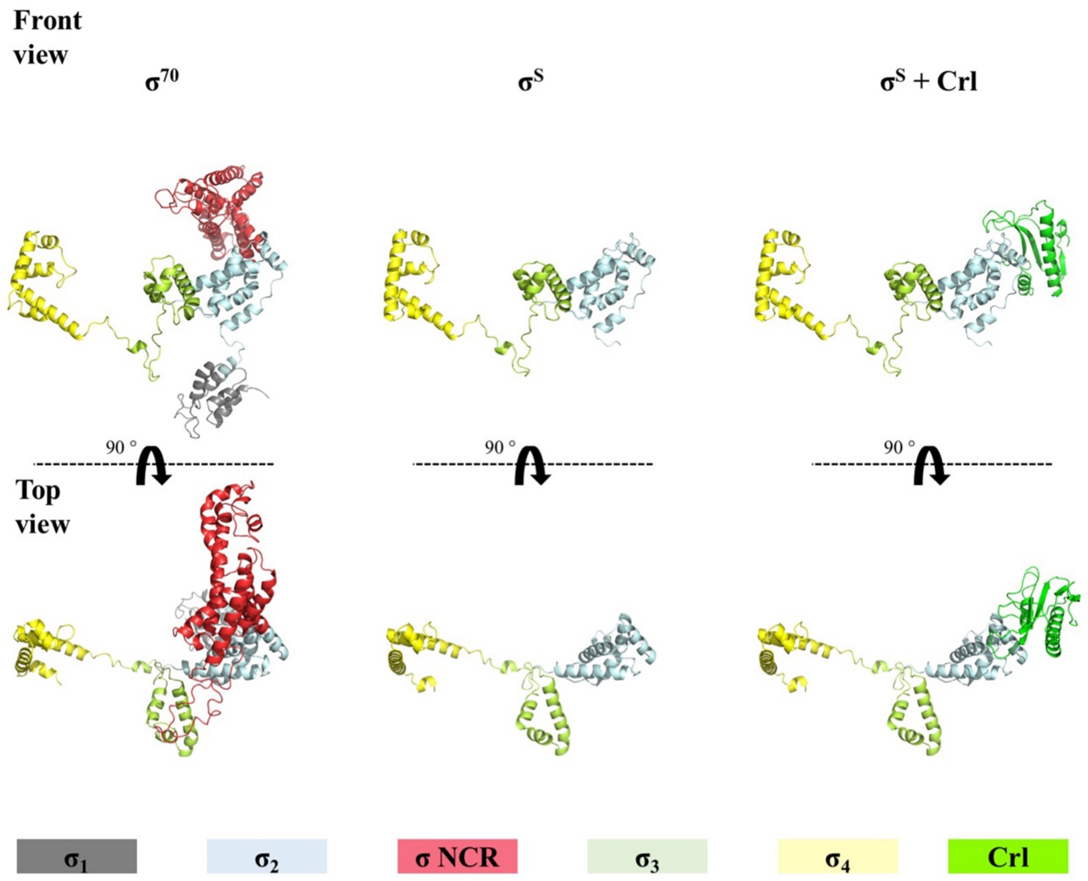

Cartagena et al., Figure S5

**Fig. S5.** Comparison of *Sty* $\sigma^{70}$ and $\sigma^{S}$ structural domain architecture.

The *Sty* $\sigma^{70}$ model was generated by homology modeling using the *Sty* $\sigma^{70}$ sequence and the structure of *Eco* $\sigma^{70}$ (from PDB 4LK1) (21) using SWISS-MODEL Workspace (22). *Sty* and *Eco* $\sigma^{70}$ have 97.6% sequence identity (Table S1).
